# Structural basis of QueC-family protein function in qatABCD anti-phage defense

**DOI:** 10.1101/2025.09.03.674047

**Authors:** Angela Gao, Douglas R. Wassarman, Philip J. Kranzusch

**Author notes:** These authors contributed equally to this work (Angela Gao, Douglas R. Wassarman).

## Abstract

QueC proteins are nucleoside biosynthesis enzymes required for production of the 7-deazaguanine derivative queuosine. Recently, QueC-family proteins were also shown to catalyze a deazaguanylation protein-nucleobase conjugation reaction in type IV CBASS bacterial anti-phage defense. Here we determine the structural basis of QueC-family protein function in a distinct bacterial immunity system named qatABCD. We demonstrate that the QueC-family protein QatC forms a specific complex with the immunity protein QatB and that this complex is minimally required for qatABCD defense. Crystal structures of the QatBC complex enable direct comparison of qatABCD and type IV CBASS defense and support a shared role for QueC-family proteins in targeting protein substrates for N-terminal modification. We show that the QatB unstructured N-terminus and N-terminal glycine motif are essential for qatABCD defense *in vivo*, suggesting a modification occurs analogous to CBASS deazaguanylation. These findings highlight broad roles of QueC proteins beyond nucleoside biosynthesis and suggest that adaptation of QueC-like proteins for specialized biochemical functions is a common strategy in bacterial anti-phage immunity.

## Introduction

Queuosine (Q) is a conserved guanosine derivative incorporated at the wobble position of Tyr, Asn, Asp, and His tRNAs in most organisms, where it enhances translation fidelity and efficiency^1–4^. Whereas plants and animals salvage Q from the environment^5,6^, bacteria synthesize Q *de novo* from GTP through a multi-enzyme biosynthetic pathway. A well characterized example of Q biosynthesis is the *Bacillus subtilis queCDEF* operon (formerly *ykvJKLM*), which encodes four enzymes required to synthesize to the key nucleobase precursor preQ17–10. Within this pathway, QueC (7-cyano-7-deazaguanine synthase) catalyzes two sequential ATP-dependent reactions that transform 7-carboxy-7-deazaguanine (CDG) into the nitrile product preQ_0_.

Enzymes related to QueC have more recently been implicated in functions beyond tRNA modification^11,12^. A defining example is Cap9, a QueC homolog encoded by the bacterial immune system type IV CBASS (cyclic oligonucleotide-based anti-phage signaling systems)^13^. Instead of producing a free nucleobase, Cap9 conjugates CDG directly to the N-terminal glycine of a protein substrate, generating a specialized deazaguanylation protein modification termed NDG (N-terminal 7-deazaguanine)^14^. NDG modification is essential for CBASS-mediated defense against phage infection, highlighting how QueC protein domains can be repurposed for distinct functions in the context of immunity. In addition to CBASS immunity, QueC domains are encoded in diverse anti-phage defense operons including the QueC-like protein QatC in qatABCD immunity^15^, but the function of QueC-family proteins in other anti-phage defense systems remains unknown.

Here we determine the structural basis of QatC function in qatABCD anti-phage defense. QatC interacts with QatB, a protein of previously unknown function, to form a complex that is minimally required for qatABCD system activity. Crystal structures of the QatBC complex reveal that QatC retains a conserved QueC core domain with distinct N-terminal and C-terminal extensions. Mirroring the requirements of NDG modification in type IV CBASS immunity, QatB contains an unstructured N-terminus and a terminal glycine motif that, along with conserved QatC active site residues, are required for anti-phage defense. Together, our findings support that QatC is a functional analog of Cap9 capable of catalyzing protein modification and demonstrate that adaptation of QueC-like proteins for specialized biochemical functions is a common strategy in bacterial anti-phage defense.

## Results

### A QatB–QatC complex is minimally required for qatACBD anti-phage defense

To define the broader function of QueC-family proteins in bacterial immunity, we compared QueC-domain-containing proteins in qatABCD and type IV CBASS anti-phage defense operons. qatABCD systems encode a QueC-domain-containing protein named QatC that is typically ∼20 kDa larger than both canonical QueC proteins and the CBASS QueC-domain containing protein Cap9 (Fig. 1a)^15^. To evaluate qatABCD anti-phage defense, we cloned and expressed a *Pseudomonas aeruginosa* qatABCD operon in *Escherichia coli* and challenged with a panel of 13 phages representing four viral families^16^. Expression of qatABCD led to reduced plaque number or size for *Siphoviridae* phages λ_vir_ and Bas25 and *Myoviridae* phages Bas39, T4, and P1_vir_ but not for the tested *Demerecviridae* or *Autographiviridae* phages (Fig. 1b).

**Figure 1.**
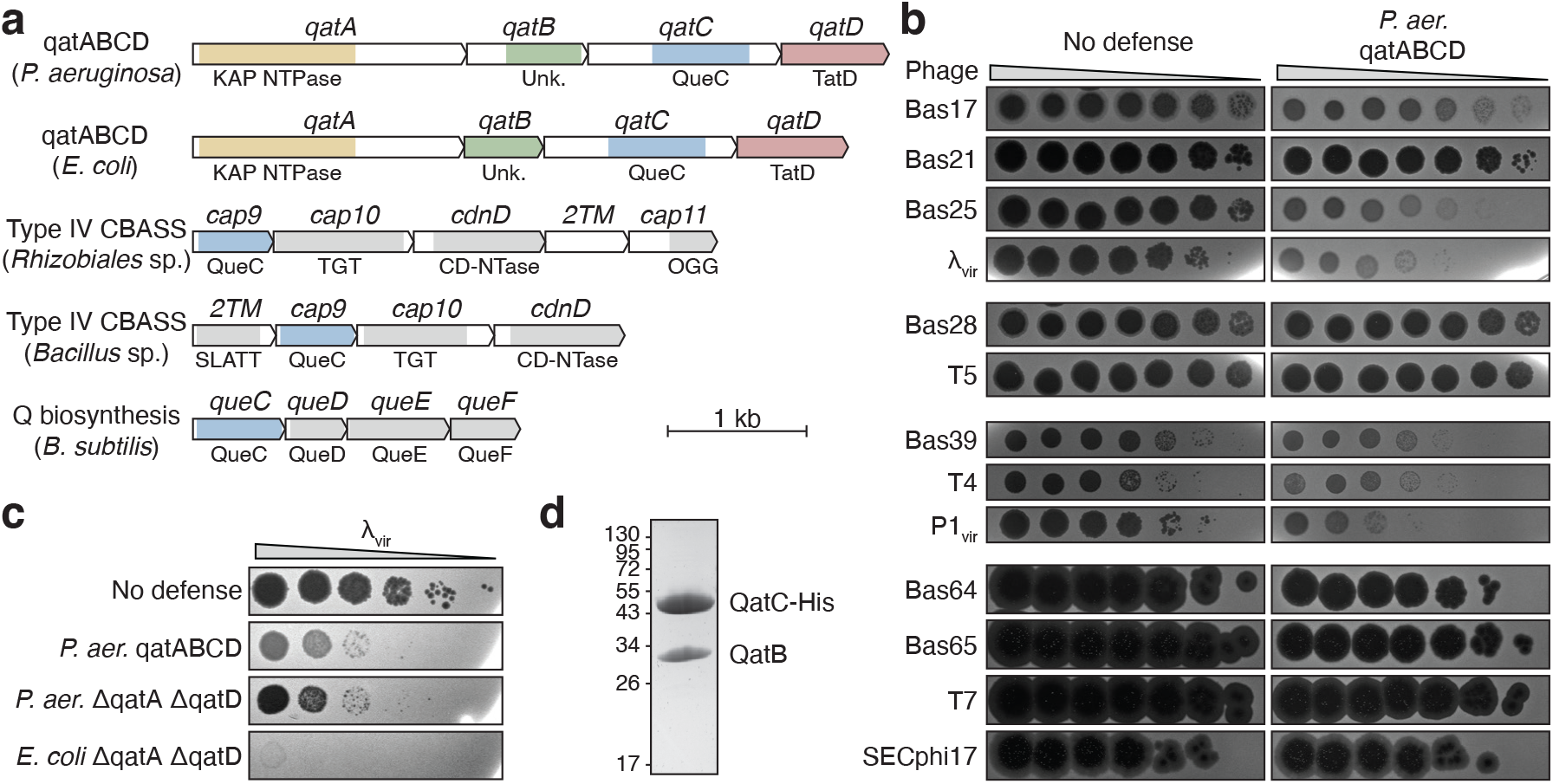
QatBC is sufficient for phage defense. **a**, Genomic architecture of qatABCD, type IV CBASS, and Q biosynthesis operons. See Supplementary Table 1 for genomic accession numbers. **b**, Representative plaque assays of *E. coli* expressing a GFP control (no defense) or a *P. aeruginosa* qatABCD operon (n=3). Phages are grouped by family (*Siphoviridae, Demerecviridae, Myoviridae, Autographiviridae*). **c**, Representative plaque assays of *E. coli* expressing the indicated construct (n=3). **d**, Coomassie-stained SDS-PAGE gel of QatBC complex purified from cells co-expressing QatB and C-terminal His-tagged QatC. Calculated molecular weights: QatB, 29.7 kDa; QatC-His, 51.6 kDa.

Three other proteins are encoded by qatABCD operons in addition to QatC: QatA, a predicted KAP-family NTPase protein; QatB, a protein of unknown function; and QatD, a predicted TatD-family nuclease protein (Fig. 1a)^15^. In addition to the KAP NTPase domain, QatA contains a C-terminal domain of unknown homology (Fig. 1a). We found that QatA is capable of multimerization and binds to both single-stranded and double-stranded DNA, though it is not predicted to have obvious dimerization or DNA-binding interfaces (Supplementary Fig. 1). Neither mutation nor deletion of QatA fully disrupted phage defense, nor did mutation or deletion of QatD (Supplementary Fig. 1).

Surprisingly, we observed that a truncated qatABCD operon lacking the *qatA* and *qatD* genes retained full defense activity against phage λ_vir_, demonstrating that QatB and QatC are alone sufficient to defend against some phages (Fig. 1c). A truncated *E. coli* qatABCD operon encoding only QatB and QatC also strongly protected against phage λ_vir_ infection (Fig. 1c). Because binding to a protein substrate is an important aspect of Cap9 function in CBASS immunity^14^, we hypothesized that QatC may form a similar protein-protein interaction with QatB. To test this, we performed pull-down experiments with His-tagged QatC and observed co-purification with untagged QatB, demonstrating that the two proteins form a stable complex (Fig. 1d). Together, these results show that QatB and QatC assemble into a protein complex that is sufficient for anti-phage defense.

### QatC is a QueC homolog adapted for qatABCD-mediated phage defense

We determined a 1.3 Å X-ray crystal structure of the *Pseudomonas aeruginosa* QatBC complex (Fig. 2a). QatB and QatC form a compact 1:1 protein assembly that exhibits both shared and distinct features compared to the 2:2 assembly formed in CBASS immunity between Cap9 and its CD-NTase protein substrate^14^. QatB is a globular alpha-helical domain protein with ten α-helices and an unstructured N-terminal region that is unresolved in the electron density (Fig. 2a; Supplementary Fig. 2). No structural homology exists between QatB and the CD-NTase protein substrate in CBASS immunity^14^. The QatC protein contains a core QueC-family Rossman-like fold composed of five parallel β-strands β5–β9 and neighboring α-helices α4–α11 that is shared with Cap9 and canonical *B. subtilis* QueC structures (Fig. 2a; Supplementary Fig. 3). Four conserved QatC cysteines C334, C354, C357, and C360 cluster to form a Zn^2+^-binding site that completes the conserved QueC-like fold (Supplementary Fig. 3). However, QatC additionally possesses N- and C-terminal extensions that are not present in Cap9 or *B. subtilis* QueC (Fig. 2b). We tested deletion constructs removing these regions and observed that they are strictly required for qatABCD anti-phage defense (Fig. 2c).

**Figure 2.**
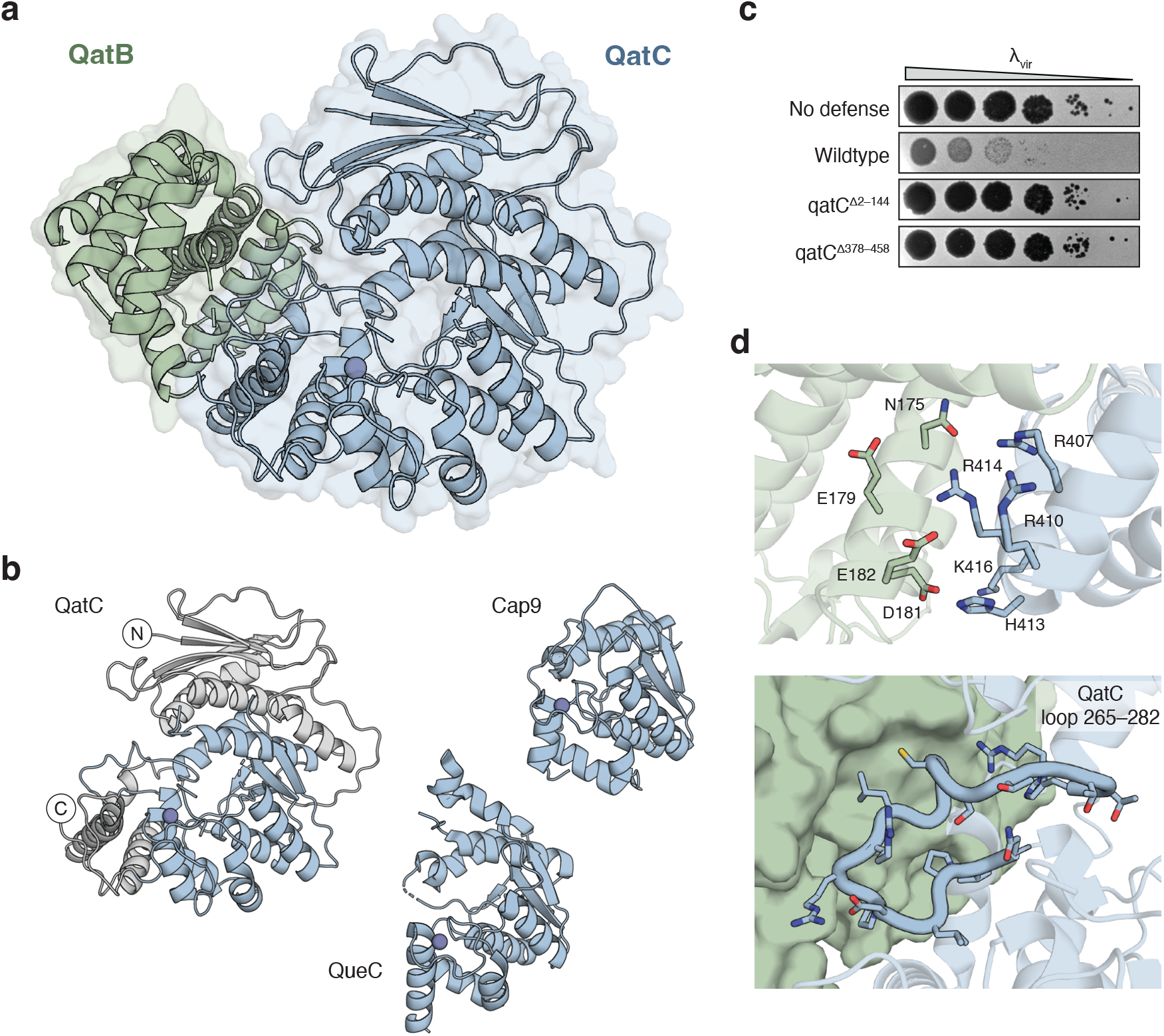
QatC interacts with QatB through unique surfaces. **a**, 1.3 Å X-ray crystal structure of the QatB-QatC apo complex. See Supplementary Table 2 for crystallographic statistics. **b**, Comparison of QatC, Cap9, and QueC structures. Shared QueC domain is highlighted in blue and QatC extensions are in grey. **c**, Representative plaque assays of *E. coli* expressing GFP control (no defense) or a *P. aeruginosa* qatABCD operon with the indicated mutations (n=3). **d**, QatB-QatC interaction interfaces. Top, polar side chains mediate interactions between a negatively charged patch on QatB (green) and a positively charged patch on the QatC C-terminal extension (blue). Bottom, QatC loop insertion in the QueC core domain (blue) makes contacts with QatB (green, surface).

The N-terminal QatC extension is composed of four parallel β-strands β1–β4 flanked by three α-helices α1–α3. The C-terminal extension consists of three α-helices α12–α14 that form the base of a large ∼1,200 Å^2^ interface with QatB (Fig. 2b). Interactions occur between a negatively charged patch of QatB centered near residue E179 and a positively charged patch of QatC near residue R410 (Fic. 2d). Additionally, QatC residues N265–H282 form a long loop extending from the QatC active site that mediates hydrophobic packing against QatB helices α7 and α8 and complementary hydrogen-bonding contacts with QatB residues Q171 and Q234 (Fig. 2d). This QatC loop bridges helices α7 and α8 of the core QueC domain and is not present in Cap9 or canonical QueC protein homologs (Supplementary Fig. 3). Each part of the QatBC interface is highly conserved (Supplementary Figs. 2 and 3), supporting that QatBC complex formation is a key feature of qatABCD defense systems.

### Conserved QatC catalytic residues are required for qatABCD anti-phage defense

QueC enzymes, including the CBASS protein Cap9, catalyze ATP-dependent reactions on a CDG (7-carboxy-7-deazaguanine) substrate molecule^10^. To evaluate whether the QatC active site may perform a similar catalytic reaction, we determined a 1.7 Å X-ray crystal structure of the QatBC complex in the presence of CDG and ATP. Although CDG was not present in the resulting structure, we observed clear density for ATP in the QatC active site, revealing the structure of a QueC enzyme in which the entire ATP molecule is resolved. Binding of ATP had minimal impact on the conformation of the QatBC complex compared to the apo complex (RMSD 1.0 Å) (Fig. 3a).

**Figure 3.**
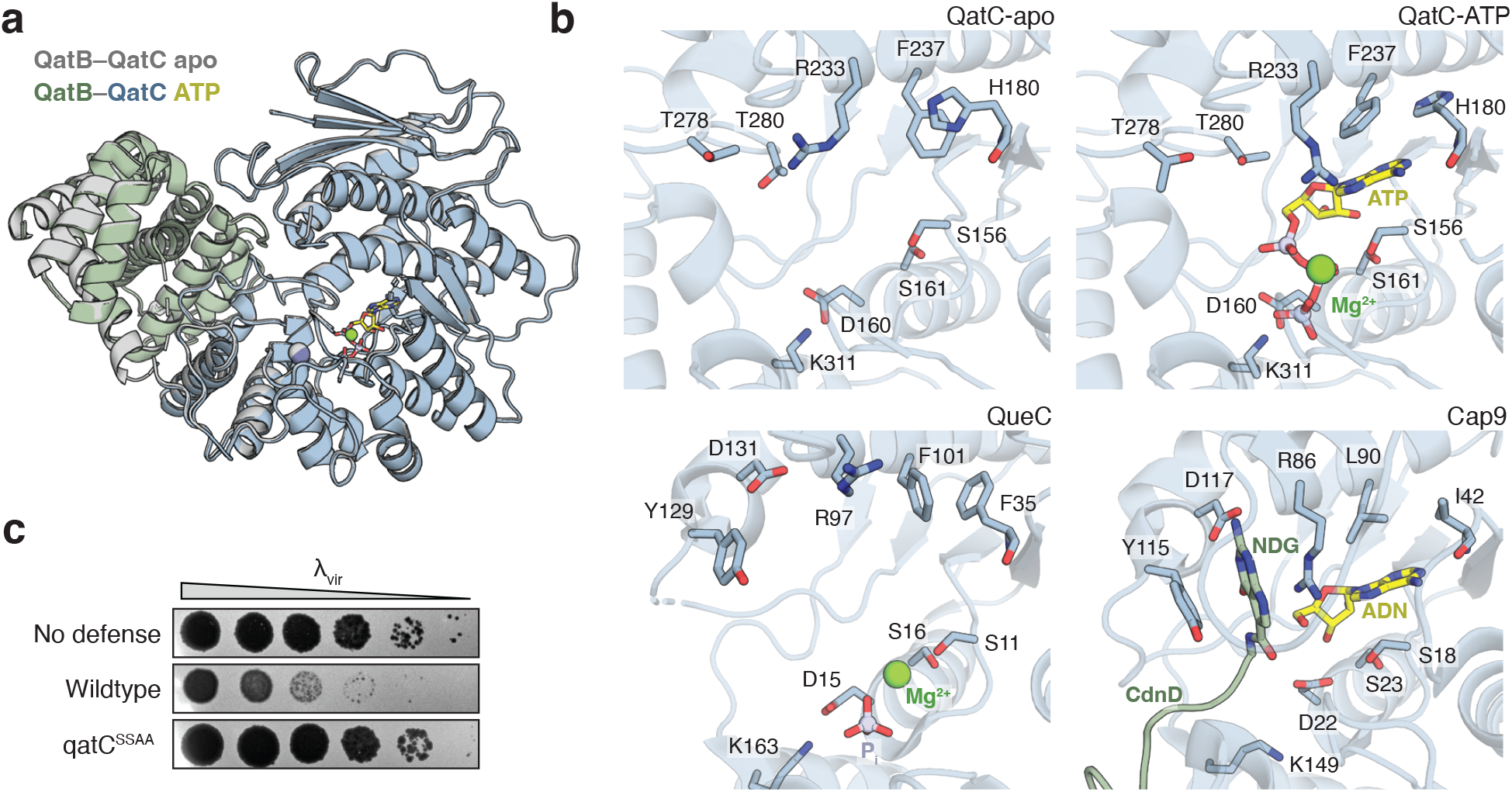
Structural analysis of the QatC active site. **a**, Comparison of the QatB-QatC apo (grey) complex with a 1.7 Å X-ray crystal structure of the QatB-QatC (green, blue) complex bound to ATP (yellow). See Supplementary Table 2 for crystallographic statistics. **b**, Comparison of QatC apo, QatC ATP-bound, QueC, and Cap9 active sites with key substrate-binding residues highlighted. Mg^2+^ ions (green), ATP or adenosine (ADN) (yellow), CdnD protein substrate with conjugated N-terminal 7-amido-7-deazaguanine (NDG) (green). PDB: 3BL5 (QueC), 9NTO (Cap9–CdnD). **c**, Representative plaque assays of *E. coli* expressing GFP control (no defense) or *P. aeruginosa* qatABCD operon (n=3). QatC S156A/S161A (qatC^SSAA^).

Analysis of the QatC active site reveals key ATP co-factor-binding and catalytic active site residues that are conserved between QueC proteins in nucleoside biosynthesis and bacterial immunity (Fig. 3b). Residues S156, D160, and S161 form a pyrophosphate-binding motif that coordinates the β- and γ-phosphates in the ATP-bound structure. The side chains of QatC residues R233 and F237 rearrange between the apo and ATP-bound structures to stabilize adenosine binding via hydrogen bond interactions around the nucleobase Hoogsteen edge and hydrophobic stacking over the aromatic rings, while the main chain of residue H180 reads out the Watson-Crick edge. The geometry of these residues that form the QatC ATP-binding pocket is conserved in structures of both *B. subtilis* QueC and Cap9.

We next analyzed the potential QatC substrate binding site. In the post-reaction structure of CBASS Cap9, the NDG nucleobase modification is coordinated by nearby Tyr and Asp residues that aromatic stack on the 7-deazaguanine nucleobase and recognize its Watson-Crick edge^14^. Analogous Tyr and Asp residues are shared in the *B. subtilis* QueC active site, supporting a highly conserved interface required for CDG substrate recognition (Supplementary Fig. 3). Interestingly, these residues are replaced in QatC by two invariant threonine residues T278 and T280 that are present in a similar arrangement in both the QatBC apo and QatBC ATP-bound structures (Supplementary Fig. 3). Alteration of the CDG substrate recognition motif in QatC suggests that the active site may bind a substrate distinct from CBASS Cap9 and canonical QueC proteins (Fig. 3b). We mutated the QatC active site residues S156A/S161A in the ATP co-factor pyrophosphate-binding-site and observed complete loss of anti-phage defense, verifying the importance of QatC catalytic function (Fig. 3c). However, despite significant effort, we were unable to identify conditions sufficient to reconstitute QatC catalytic activity *in vitro* or determine the complete catalytic function of QatC during anti-phage defense. Together these results demonstrate that QatC catalytic activity is an essential aspect of immune function but that QatC catalyzes a yet unresolved enzymatic reaction.

### QatB contains a flexible N-terminal tail with an essential glycine motif

We next used the QatBC structure to analyze potential essential features of the uncharacterized immunity protein QatB. Structural homology searches of QatB using FoldSeek^17^ and DALI^18^ revealed no significant matches to any proteins of known function in the Protein Data Bank or AlphaFold databases, nor does it have an obvious active site or substrate-binding pocket. A structure-guided alignment of QatB proteins from across qatABCD operons demonstrates that the core alpha-helical domain fold is conserved, as well as the presence of an N-terminal region predicted to be disordered^19,20^ (Supplementary Fig. 2). The N-terminal region in *P. aeruginosa* QatB is 65 amino acids long, though the first 63 residues are unresolved in the electron densities of both the apo and ATP-bound structures (Fig. 4a). Many QatB proteins are predicted to possess a similarly large N-terminal extension ranging within 30-90 amino acids long (Supplementary Fig. 2). However, in some QatB sequences, the N-terminal tail and helices α1–α3 are absent entirely. Manual inspection of a subset of such sequences suggests that in most cases this is due to misannotation of the QatB start site.

**Figure 4.**
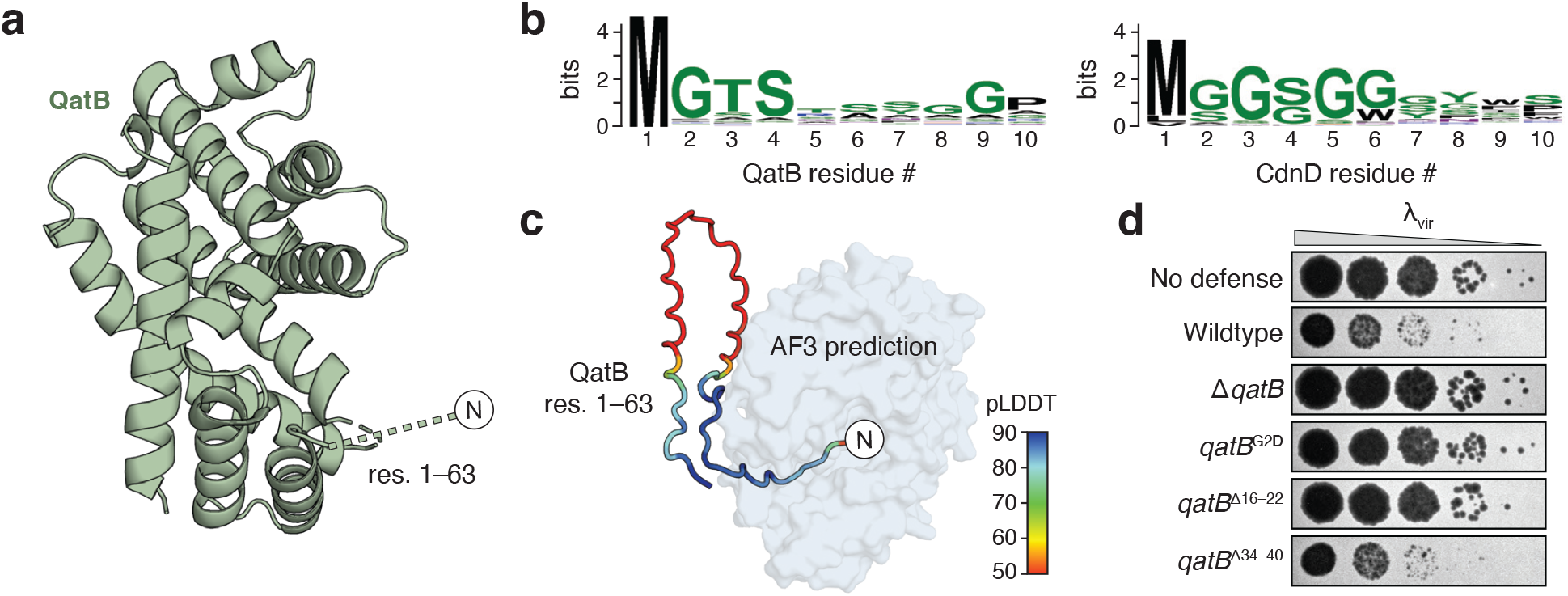
The QatB N-terminal tail is essential for phage defense. **a**, QatB structure highlighting the unresolved N-terminal tail. **b**, Sequence logos showing conservation of QatB and CdnD N-termini. **c**, Conformation of unresolved QatB tail predicted by AlphaFold 3 overlaid on the QatBC crystal structure. QatB residues colored by calculated pLDDT. **d**, Representative plaque assays of *E. coli* expressing GFP control (no defense) or *P. aeruginosa* qatABCD operon with the indicated mutations (n=3).

Notably, the alignment of QatB proteins reveals a highly conserved MGTS motif at the extreme N-terminus similar to the CD-NTase N-terminal MGGS motif required for NDG modification in type IV CBASS immunity (Fig. 4b). In bacteria, the initiating methionine is typically removed when followed by a small residue^21^. In the case of type IV CBASS, this results in an exposed glycine G2 that is post-translationally modified with NDG by Cap9^14^. We modeled the QatBC complex using AlphaFold 3 with a truncated QatB sequence where the N-terminal methionine was removed and observed that in all models, the QatB N-terminal residues were placed inside the QatC active site in an interaction mirroring positioning of the CD-NTase N-terminus during Cap9-mediated deazaguanylation (Fig. 4c)^14,22^.

These experimental and modeling results support that QatB may serve as a protein substrate of the QatC QueC domain in a reaction mechanistically shared between qatABCD and type IV CBASS anti-phage defense. To further test this model, we mutated the QatB N-terminus and measured the impact on qatABCD function *in vivo*. In CBASS, substitution of the G2 residue with a bulky, charged aspartate prevents deazaguanylation and blocks defense^14^. Likewise, a qatABCD operon encoding the QatB G2D mutant exhibited no phage defense activity (Fig. 4d). Additional conserved residues are found along the beginning of the QatB tail, particularly at Pro16 and Trp18. Deletion of QatB residues 16-22 containing these conserved positions resulted in loss of anti-phage defense, while deletion of a stretch of relatively variable residues 34–40 had no impact (Fig. 4d; Supplementary Fig. 2). Together, these results define functional roles for QatBC complex formation in qatABCD anti-phage defense and suggest that the protein deazaguanylation reaction observed in type IV CBASS is a shared mechanism of anti-phage defense.

## Discussion

Here we define the QatBC protein complex as the core functional unit of the qatABCD anti-phage defense system and determine structures that explain how the QueC-family protein QatC contributes to bacterial immunity. Our findings extend recent work on type IV CBASS^14^, demonstrating that QueC-family proteins act as key catalytic components across multiple defense systems. In contrast to CBASS, which requires several proteins and activation of a signaling pathway for defense, a minimal protein assembly consisting of QatC and its putative substrate QatB is sufficient for qatABCD anti-phage activity. Together, these findings expand the known roles of QueC-family proteins beyond nucleoside biosynthesis.

Crystal structures of the QatBC complex reveal a series of catalytic and protein scaffolding features required for qatABCD defense. QatC exhibits a canonical QueC fold with conserved architectural elements and catalytic residues. Distinct QatC-specific extensions beyond the core QueC domain form a binding interface with QatB and are required for anti-phage defense (Figs. 2 and 3). QatB, a protein of previously unknown function, contains a helical domain and a long, unstructured N-terminal region bearing a highly conserved N-terminal MGTS motif. Mutational analyses confirm that both QatC catalytic activity and the QatB N-terminal MGTS motif are essential for qatABCD immune function *in vivo* (Figs. 2–4)

These results raise the central mechanistic question of what reaction QatC catalyzes in qatABCD immunity. Conservation of the active site geometry and the dependence on the N-terminal glycine of QatB suggest a protein modification analogous to the NDG modification found in type IV CBASS^14^. Notably, QatB possesses all the essential features of an NDG substrate, including complex formation with a QueC protein, positioning of a long, unstructured N-terminus in proximity of the QueC active site, and an essential N-terminal glycine motif that that serves as the acceptor site for NDG modification in CBASS. These findings support a shared mechanism in bacterial immunity where QueC-family enzymes target a partner protein for modification (Fig. 5). However, the precise chemical reaction catalyzed by QatC and how modification of QatB leads to restriction of phage infection remain key open questions for future studies. QatC diverges from Cap9 and other QueC proteins by substituting invariant threonine residues for the canonical Tyr/Asp residues that recognize nucleobase substrate (Fig. 3b), which may suggest a distinct protein modification is produced. Further divergence of QatC is evidenced by its extension domains, which are strictly required for defense but lack obvious catalytic motifs. Part of the C-terminal extension participates in QatB binding, but the remaining regions may play additional regulatory roles. These differences may explain why QatC activity has been difficult to reconstitute *in vitro*, pointing to a requirement for additional cofactors, conformational control, or infection-dependent activators.

**Figure 5.**
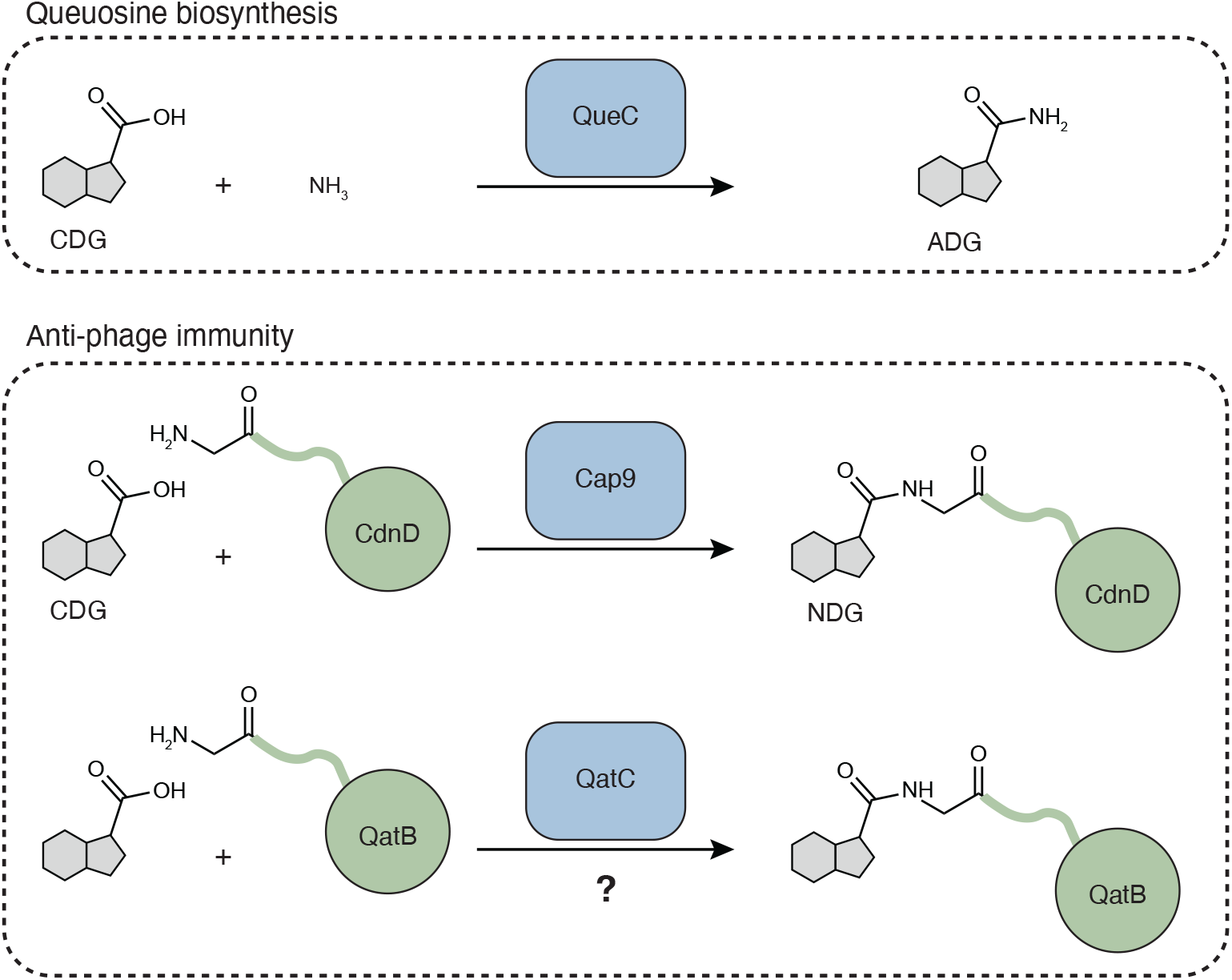
Model of possible QatBC complex function. Top, first step of the QueC-mediated nucleobase synthesis reaction. QueC catalyzes activation of the 7-carboxy-7-deazaguanine (CDG) carboxylic acid using a molecule of ATP, followed by nucleophilic attack with NH_3_ to generate 7-amido-7-deazaguanine (ADG). Middle, Cap9 catalyzes an analogous reaction with CDG using the N-terminal amine of CdnD as the nucleophile in place of NH_3_ to form an N-terminal 7-deazaguanine (NDG) protein modification. Bottom, proposed parallel model of QatC-mediated modification of QatB N-terminus.

Finally, our discovery that the QatBC complex alone confers defense against phage infection leads to new questions about the roles of QatA and QatD. The annotation of QatA as an NTPase and QatD as a nuclease led to the suggestion that these proteins are effectors^15^, but mutation or deletion of QatA and QatD has limited impact on phage defense. These enzymes may contribute to regulation of the central QatBC immune complex, and further study may explain why they are included in nearly all qatABCD operons. Although the full pathway of qatABCD-mediated defense remains unknown, our results demonstrate that qatABCD utilizes a mechanism that is both similar to and distinct from type IV CBASS and introduces a modular family of phage defense systems that rely on a shared QueC-derived reaction.

## Methods

### Bacterial strains and phages

This study used *E. coli* strains Top10, BL21(DE3) RIL, and BW25113. Top10 and BL21(DE3) RIL cells were grown in lysogeny broth (LB). BW25113 cells were grown in LB supplemented with 5 mM MgCl_2_, 5 mM CaCl_2_, 0.1 mM MnCl_2_ (MMC). Bacterial plates were prepared with 1.5% agar and top agar was prepared with 0.5% low-melt agar. When applicable, media and plates were supplemented with ampicillin (100 µg mL^−1^) or chloramphenicol (34 µg mL^−1^) to maintain plasmids.

Phages were propagated by picking a single plaque into SM buffer (50 mM Tris-HCl pH 7.5, 100 mM NaCl, 8 mM MgSO_4_), followed by inoculation into a liquid bacterial culture grown to optical density at 600 nm (OD600) of 0.3. Cultures were incubated while shaking at 37°C until culture collapse, then centrifuged for 10 min at 4000 × g. The supernatant was filtered through a 0.2 µm filter and stored at ^−^4°C.

### Plasmid construction

Native qatABCD operon sequences were synthesized and cloned into an arabinose-inducible pBAD vector containing an ampicillin resistance cassette (Twist Bioscience). For protein expression, sequences were subcloned into an IPTG-inducible pET vector containing an ampicillin resistance cassette via PCR and Gibson assembly. Mutations, deletions, and 6×His tags were introduced by site-directed mutagenesis PCR. A ribosome binding site was inserted when deleting genes that overlapped with a subsequent coding sequence to prevent disruption of translation. Plasmids were transformed into chemically competent Top10 cells for propagation. All plasmids were confirmed by whole plasmid sequencing (Plasmidsaurus).

### Phage plaque assays

Constructs for plaque assays consisted of the qatABCD operon cloned into the pBAD vector with any indicated mutations or deletions. The ΔqatA ΔqatD construct additionally contained a 6×His tag on the C-terminus of QatC. sfGFP cloned into the pBAD vector was used as a no defense control. Plasmids were transformed into chemically competent BW25113 cells and liquid cultures were inoculated from single colonies or glycerol stocks. Cultures were incubated while shaking at 37°C until they reached an OD600 of at least 0.3. Top agar was prepared by diluting cultures to an OD600 of 0.06 and supplemented with 0.2% L-arabinose, then poured onto LB plates and allowed to solidify at room temperature for 1 hour. 6 mL of top agar was used for 100×15mm round plates and 15 mL was used for 120×120mm square plates. A 10-fold dilution series was prepared for each phage in SM buffer and 2.5 µL of each dilution was spotted onto solidified plates. Drops were dried at room temperature and plates were incubated overnight at 30°C. Plates were imaged using a ChemiDoc MP Imaging System (Bio-Rad).

### Protein expression and purification

All proteins were expressed from a custom pET vector^23^. QatC was C-terminally tagged with a 6×His tag and co-expressed with QatB. QatA was expressed with a C-terminal 6×His tag alone or within the full qatABCD operon. Protein constructs were transformed into chemically competent BL21(DE3) RIL cells. and plated on 1.5% agar prepared with MDG media (2 mM MgSO_4_, 0.5% glucose, 0.25% aspartic acid, 25 mM Na_2_HPO_4_, 25 mM KH_2_PO_4_, 50 mM NH_4_Cl, 0.5 mM Na_2_SO_4_, 2–50 µM trace metals). Three to four colonies were inoculated into 30 mL of MDG liquid culture and incubated while shaking at 37°C overnight. 12 mL of overnight culture was inoculated into 1 L of M9ZB (47.8 mM Na_2_HPO_4_, 22 mM KH_2_PO_4_, 18.7 mM NH_4_Cl, 85.6 mM NaCl, 1% casamino acids, 0.5% glycerol, 2 mM MgSO_4_, 2–50 µM trace metals) and incubated while shaking at 37°C until cultures reached an OD600 of 2.5–3.0. Protein expression was induced with 0.5 mM isopropyl-β-d-thiogalactoside (IPTG), and cultures were incubated while shaking at 16°C overnight. Cell pellets were harvested by centrifugation, washed with 1× PBS, and lysed by sonication in lysis buffer (20 mM HEPES-KOH pH 7.5, 400 mM NaCl, 10% glycerol, 30 mM imidazole, 1 mM DTT). Lysates were clarified by centrifugation at 40,000 × g for 30 minutes and the supernatant was poured over 8 mL of Ni-NTA resin (Qiagen) equilibrated in lysis buffer. The resin was washed twice with lysis buffer, or with lysis buffer and wash buffer (20 mM HEPES-KOH pH 7.5, 1 M NaCl, 10% glycerol, 30 mM imidazole, 1 mM DTT) for QatA expressed alone. Proteins were eluted with elution buffer (20 mM HEPES-KOH pH 7.5, 400 mM NaCl, 10% glycerol, 300 mM imidazole, 1 mM DTT) and concentrated using centrifugal filter units (Millipore Sigma). Samples were centrifuged at 12,000 × g for 10 minutes to remove precipitates and applied to a 16/600 Superdex 200 column (Cytiva) equilibrated in gel filtration buffer (20 mM HEPES-KOH pH 7.5, 250 mM KCl, 1 mM TCEP-KOH). Purified proteins were concentrated using centrifugal filter units (Millipore Sigma) to at least 10 mg mL^−1^. Aliquots were flash frozen in liquid nitrogen and stored at ^−^80°C.

### X-ray crystallography

Crystals were grown in 2 μL hanging drops at 18°C in EasyXtal 15-well trays (NeXtal). Crystals of the QatB-QatC apo complex were grown in a 1.2:0.8 mixture of 5 mg mL^−1^ protein in gel filtration buffer and reservoir solution consisting of 18% PEG-3350 and 0.1 M NaCl. Crystals of the ATP-bound complex were grown in a 1.2:0.8 mixture of 5 mg mL^−1^ protein in gel filtration buffer with 1 mM ATP and 1 mM CDG, and reservoir solution consisting of 14% PEG-20,000 and 1 M MES pH 6.3. Crystals were cryo-protected with reservoir solution supplemented with 22% ethylene glycol for the apo complex and 36% for the ATP-bound complex and frozen in liquid nitrogen. X-ray diffraction data for the apo complex was collected at National Synchrotron Light Source II beamline 17-ID-1 and processed using autoPROC^24^ including correction for anisotropy using STARANISO^25^. The initial search model was generated using AlphaFold 3^22^. X-ray diffraction data for the ATP-bound complex was collected at Advanced Photon Source beamline 24-ID-E and processed using RAPD2. The apo structure was used as the search model. Molecular replacement and model refinement were performed using PHENIX^26^ and model building was performed in COOT^27^. Statistics are described in Supplementary Table 1. Structure figures were generated with PyMOL (Schrödinger).

### Electrophoretic mobility shift assays

All substrates had the same core sequence (5′-ACTGCACTACAACAGAACCAGAGCGACTGCACTACAACAGAACCA-3′). Overhangs were composed of a 15 nt poly-T sequence on the forward strand. DNAs were prepared in annealing buffer (10 mM Tris-HCl pH 7.5, 50 mM NaCl) and double-stranded substrates were annealed by incubation at 95°C for 5 minutes followed by cooling to 25°C over 100 minutes. Reactions were performed with 1 μM of substrate and 0, 2 μM, 10 μM, 20 μM, 30 μM, or 50 μM QatA in 20 mM HEPES-KOH pH 7.5, 100 mM KCl, and 1 mM TCEP for 30 min at room temperature. 3 μL of 50% glycerol was added to each 20 μL reaction and 10 μL of each sample was loaded onto a 2% TB (Tris-borate) agarose gel. The gel was run at 250 V for 45 minutes, post-stained with TB buffer containing 10 μg mL^−1^ ethidium bromide for 30 minutes, and destained in water for 30 minutes. Gels were imaged using a ChemiDoc MP Imaging System (Bio-Rad).

### Bioinformatic analysis

Protein sequences were obtained from InterPro^28^. Sequence logo diagrams were generated using WebLogo 3^29^ using the first 10 amino acids of all QatB sequences. For full-length sequence alignments, QatB or QatC sequences were clustered using MMseqs2^30–32^ with minimum sequence identity of 0.5 and minimum alignment coverage of 0.8. Representative sequences were selected from the largest clusters and were aligned to QatB or QatC/QueC reference sequences using MAFFT with default parameters^31–33^. Alignments were visualized using Jalview^34^. Protein structure predictions were generated using AlphaFold 3^22^.

## Statistics and reproducibility

Experimental details regarding replicates and sample size are described in the figure legends.

## Acknowledgements

The authors are grateful to members of the Kranzusch laboratory for helpful comments and discussion. The work was funded by grants to P.J.K. from the Burroughs Wellcome Fund PATH program, The G. Harold and Leila Y. Mathers Charitable Foundation, The Mark Foundation for Cancer Research, the Cancer Research Institute, the Parker Institute for Cancer Immunotherapy, the Massachusetts Consortium on Pathogen Readiness (MassCPR), and the National Institutes of Health (1DP2GM146250-01). D.R.W. is supported by the Helen Hay Whitney Foundation.

X-ray data were collected at the Advanced Photon Source beamline 24-ID-E and Brookhaven National Laboratory beamline NSLS-II. This research was performed on APS beam time award(s) (DOI: https://doi.org/10.46936/APS-189898/60014017) from the Advanced Photon Source, a U.S. Department of Energy (DOE) Office of Science user facility operated for the DOE Office of Science by Argonne National Laboratory under Contract No. DE-AC02-06CH11357. The Center for Bio-Molecular Structure (CBMS) is primarily supported by the NIH-NIGMS through a Center Core P30 Grant (P30GM133893), and by the DOE Office of Biological and Environmental Research (KP1607011). NSLS-II is a U.S. DOE Office of Science User Facility operated under Contract No. DE-SC0012704. This publication resulted from data collected using beamtime obtained through NECAT BAG proposal #317877.

## Author Contributions

The study was designed and conceived by A.G., D.R.W. and P.J.K. Data were collected and analyzed by A.G. and D.R.W. The manuscript was written by A.G., D.R.W. and P.J.K. All authors support the conclusions.

## Additional Information

Correspondence and requests for materials should be addressed to P.J.K. All illustrations were created using Adobe Illustrator.

## Data Availability Statement

All data are available in the manuscript or the supplementary information. The crystal structures of apo and ATP-bound QatB–QatC have been deposited in the Protein Data Bank under accession codes 9Y01 and 9Y02, respectively.

## Supplementary Figures

**Supplementary Figure 1.**
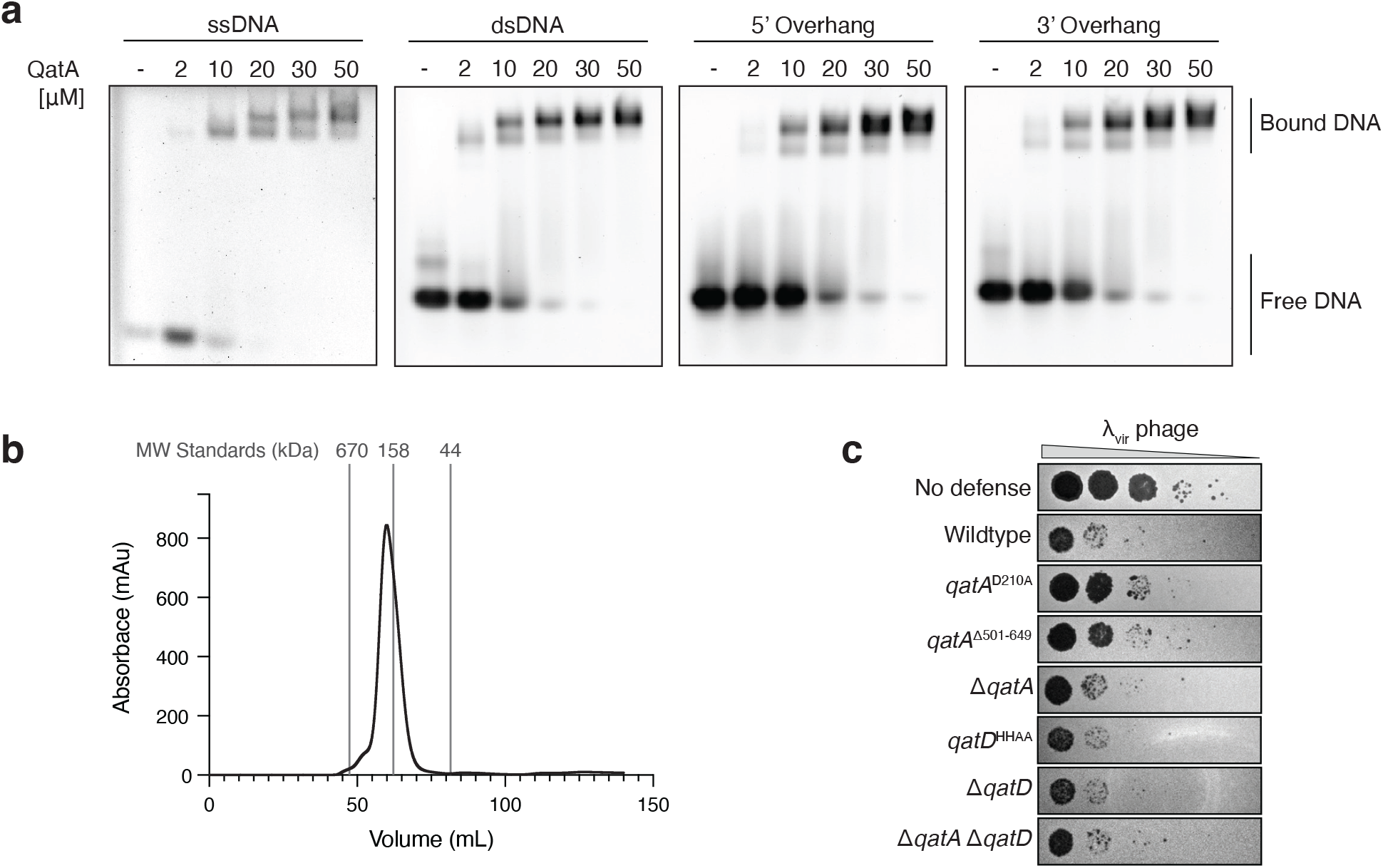
QatA and QatD in qatABCD-mediated phage defense. **a**, Electrophoretic mobility shift assays of QatA on ssDNA, dsDNA, 5′ overhang, and 3′ overhang substrates run on 2% TB-agarose gels stained with ethidium bromide. Protein concentrations are expressed as μM of QatA monomer. **b**, Size exclusion chromatography trace of His-tagged QatA expressed in the full qatABCD operon. Calculated molecular weight of QatA monomer is 72 kDa and QatA dimer is 144 kDa. Peak elution volumes of molecular weight standards are indicated by vertical grey lines. Thyroglobulin, 670 kDa; γ-globulin, 158 kDa; ovalbumin, 44 kDa. **c**, Representative plaque assays of *E. coli* expressing GFP control (no defense) or *P. aeruginosa* qatABCD with the indicated (n=4). QatD H15A/H17A (qatD^HHAA^).

**Supplementary Figure 2.**
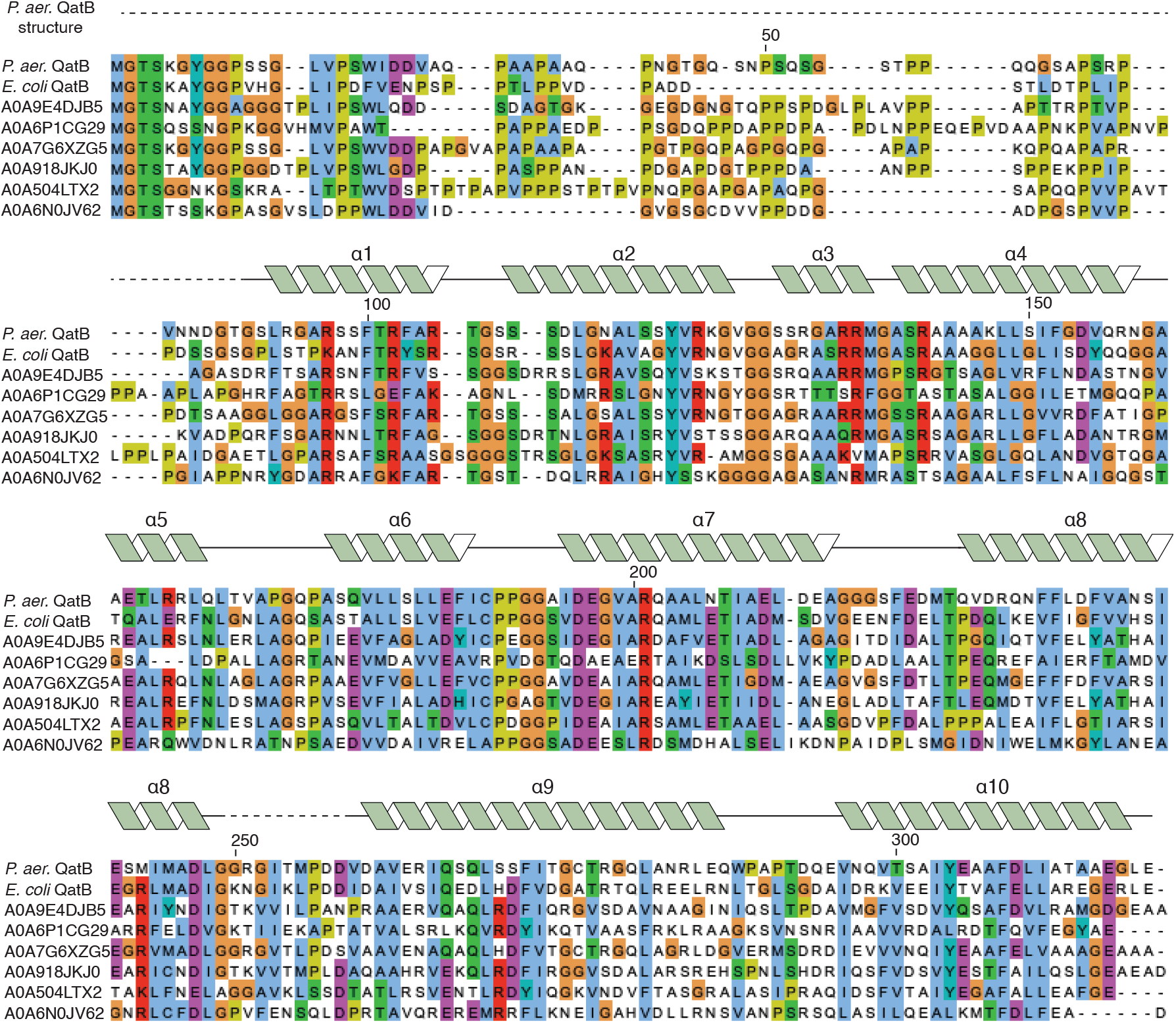
QatB alignment. Sequence alignment of representative QatB sequences (UniProtKB accession numbers shown). Secondary structure is annotated from *P. aeruginosa* QatB structure in complex with QatC. Alignment is colored using ClustalX color scheme based on side chain properties.

**Supplementary Figure 3.**
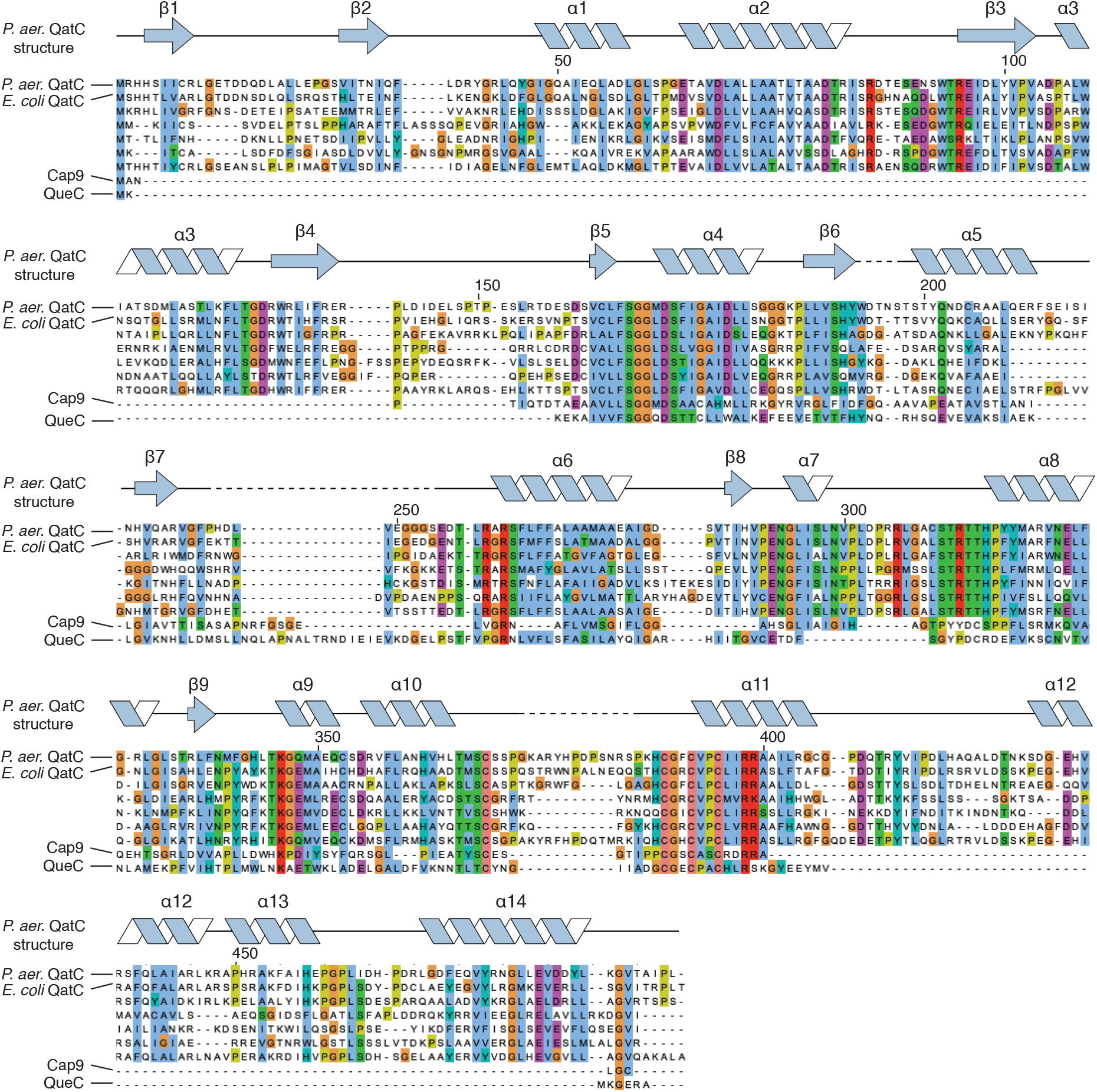
QatC alignment. Sequence alignment of representative QatC sequences (UniProtKB accession numbers: A0A2P6WBR3, A0A1E2V6V1, A0AA91FJV8, A0A158EPR1, A0A7X2LQZ7), *Rhizobiales sp*. Cap9, and *B. subtilis* QueC. Secondary structure is annotated from *P. aeruginosa* QatC structure in complex with QatB. Alignment is colored using ClustalX color scheme based on side chain properties.

